# DNAJB1-PRKACA fusion induces liver inflammation in zebrafish model of Fibrolamellar Hepatocellular Carcinoma

**DOI:** 10.1101/781583

**Authors:** Sofia de Oliveira, Ruth A. Houseright, Benjamin G. Korte, Anna Huttenlocher

## Abstract

Fibrolamellar Hepatocellular Carcinoma (FLC) is a rare liver cancer that primarily affects adolescents and young adults. Up to now there is only one identified molecular target responsible for driving the disease, the chimeric protein encoded by DNAJB1-PRKACA (J-PKAca). Immune cells have been identified as key players in liver cancer biology, however the effect of J-PKAca on inflammation in the liver microenvironment is not known. Here we report a new zebrafish model of FLC with non-invasive live imaging capabilities that allows the study of the interactions between immune cells and transformed hepatocytes. We found that overexpression of the dnajb1a-prkcaaa fusion gene specifically in hepatocytes induces early malignancy features in FLC transgenic larvae, such as increased liver and hepatocyte size. In addition, this aberrant form of PKA promotes a pro-inflammatory liver microenvironment by increasing the number of neutrophils and macrophages in the liver area and inducing macrophage polarization to a pro-inflammatory phenotype. Increased caspase-a activity was also found in the liver of FLC transgenic larvae. Importantly, pharmacological inhibition of TNFα secretion and caspase-a activity decreased liver size and inflammation. Overall, these findings suggest that inflammation may be an early feature of FLC involved in progression, and that targeting TNFαand caspase-1 may be beneficial in treating FLC.

## Introduction

Fibrolamellar Hepatocellular Carcinoma (FLC) is a rare and understudied liver cancer that primarily affects adolescents and young adults. Surgery (resection/liver transplantation) is the most common treatment with the best prognosis for FLC patients; however, recurrence is very common in FLC patients. Unfortunately, few therapeutic strategies are available, and they are not very effective. Most studies have focused on identifying molecular targets that drive the disease and specific FLC tumor markers (Dhingra et al., 2010; Graham et al., 2015; Honeyman et al., 2014; Li et al., 2010; Oikawa et al., 2015; Ross et al., 2011; Simon et al., 2015). There is only one identified molecular target responsible for driving the disease, resulting from a 400kb deletion on chromosome 19, the DNAJB1-PRKACA fusion transcript (Honeyman et al., 2014; Simon et al., 2015). Recently developed murine models using CRISPR/Cas9 or overexpression of this fusion gene support the idea that the fused DNAJB1-PRKACA gene alone is sufficient to drive tumorigenesis *in vivo (Engelholm et al., 2017; Kastenhuber et al., 2017)*.

The biochemical features of the chimeric protein encoded by DNAJB1-PRKACA are a major research focus in the field (Graham et al., 2018; Riggle et al., 2016). The DNAJB1-PRKACA transcript comprises the J-domain of DnaJB1 (the amino-terminal 69 residues) fused to the carboxyl-terminal 336 residues of the PKAcα. Importantly, the chimeric protein, J-PKAcα, is enzymatically active (Simon et al., 2015; Xu et al., 2015). However, it is still unclear how the physiological properties of this aberrant PKA cause FLC. PKA is a key regulator of both the innate and adaptive immune response (Serezani et al., 2008; Skalhegg et al., 2005). Innate immune cells play a role in liver cancer development and progression (de Oliveira et al., 2019; Kuang et al., 2009; Kuang et al., 2011; Li et al., 2015; Yan et al., 2015; Yan et al., 2017), but the effect of J-PKAcα on immune cells and subsequent modulation of the liver microenvironment and FLC progression is still unclear. There is a need for animal models amenable to studying immune cell-tumor cell interactions with non-invasive live imaging. Therefore, to study immune cell-cancer cell interactions in an intact animal *in vivo* we developed a FLC zebrafish model. Zebrafish have unmatched live imaging capabilities and scalability, and they are amenable to whole-organism-level experiments and genetic and pharmacological manipulations. Hepatocyte-specific overexpression of the *dnajb1a-prkcaa* fusion transcript promotes early malignancy features in zebrafish FLC larvae and formation of masses in adults. In addition, expression of J-PKAcα in hepatocytes induces infiltration of neutrophils and macrophages into the liver area and macrophage polarization to a pro-inflammatory phenotype in 7 days-post-fertilization FLC transgenic larvae. Caspase-a activity in the liver is also increased. Finally, using a pharmacological approach to target inflammation and FLC progression, we found that inhibition of TNFα or caspase-a decreased neutrophil and macrophage infiltration and liver size in FLC larvae. Overall, our data suggest that inflammation occurs early in FLC larvae and that pharmacological inhibition of TNFα secretion and caspase-a activity might be targets to treat inflammation and progression in FLC.

## Results

### Overexpression of dnajb1a-prkacaa in hepatocytes can induce tumorigenesis in adult zebrafish

The DNAJB1-PRKACA chimera is a unique marker for FLC. This chimera is a fusion product of a 400Kb deletion on chromosome 19 that occurs in the liver of FLC patients (Fig. 1A). Expression of the DNAJB1-PRKACA fusion transcript is sufficient to drive tumorigenesis *in vivo* in murine models *(Engelholm et al., 2017; Kastenhuber et al., 2017)*. Zebrafish has two homologous genes for *dnajb1, dnajb1a* (ENSDARG00000099383) and *dnajb1b* (ENSDARG00000041394), located in chromosomes 3 and 1 respectively. In addition, there are also two homologous genes for *prkaca*, *prkacaa* (ENSDARG00000100349) and *prkacab* (ENSDARG00000016809), also located in chromosomes 3 and 1 respectively. Here, using the hepatocyte-specific *fabp10a* promoter, we overexpressed a zebrafish *dnajb1a-prkacaa* chimera, with 91.6% identity and 97% similarity with its human counterpart (Fig. 1B). Using the transposase system, we generated a stable line, referred to as *Tg(fabp10a:dnajb1a-prkacaa_cryaa:Cerullean).* This was generated in the pigment-deficient Casper background to enable non-invasive live imaging at later developmental stages. To facilitate liver visualization, we outcrossed the FLC line to a transgenic line expressing egfp-l10a, *Tg(fabp10a:egfp-l10)* (Table 1) (Fig. 2A). To determine if overexpression of *dnajb1a-prkacaa* fusion transcript was able to induce tumorigenesis in zebrafish, we dissected livers from 8 and 12-month-old FLC and control fish and performed a blinded, conventional histopathological evaluation of hematoxylin and eosin-stained sections. Compared with controls, FLC livers are larger and display mildly disrupted hepatocellular architecture, characterized by increased thickness of hepatocellular cords. Hepatocytes from FLC livers also have vesiculated chromatin and prominent and sometimes multiple nucleoli (Fig.2 C). In 1/8 fish at 8 months and an additional 1/7 fish at 12 months (Fig. 2B), unencapsulated masses are noted within the hepatic parenchyma. Masses lack typical hepatic architecture and consist of disorganized sheets of well-differentiated hepatocytes traversed by blood vessels (Fig. 2C). Staining with Masson’s trichrome yielded no difference in collagen deposition between FLC livers and controls (data not shown). Our findings suggest that *dnajb1a-prkacaa* overexpression specifically in hepatocytes can induce features associated with tumorigenesis *in vivo* in zebrafish.

**Figure 1:**
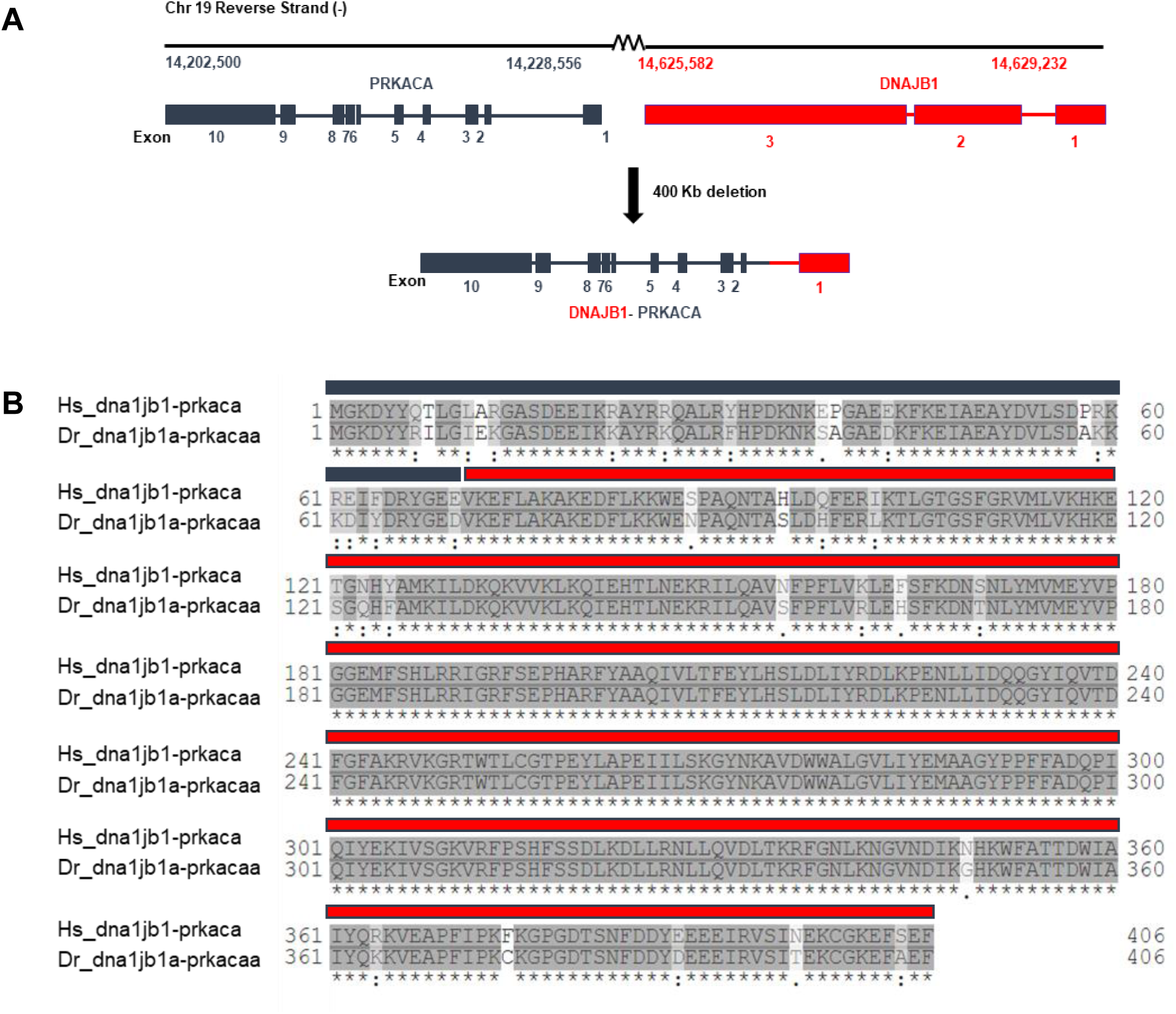
Schematic of DNAJB1-PRKACA chimera in FLC. **(A)** Chromosomic position, Exon/Intron diagram of *dnajb1* and *prkaca* genes, and fusion product after heterogenic deletion of 400Kb observed in FLC patients; **(B)** Clustal Omega alignment of *human (Hs) and zebrafish (Dr)* aa sequences corresponding to Exon 1 of Dnajb1a (blue) and to Exon 2-10 of Prkacaa (red) (Identity: 91.6%, Similarity: 97%).

**Figure 2:**
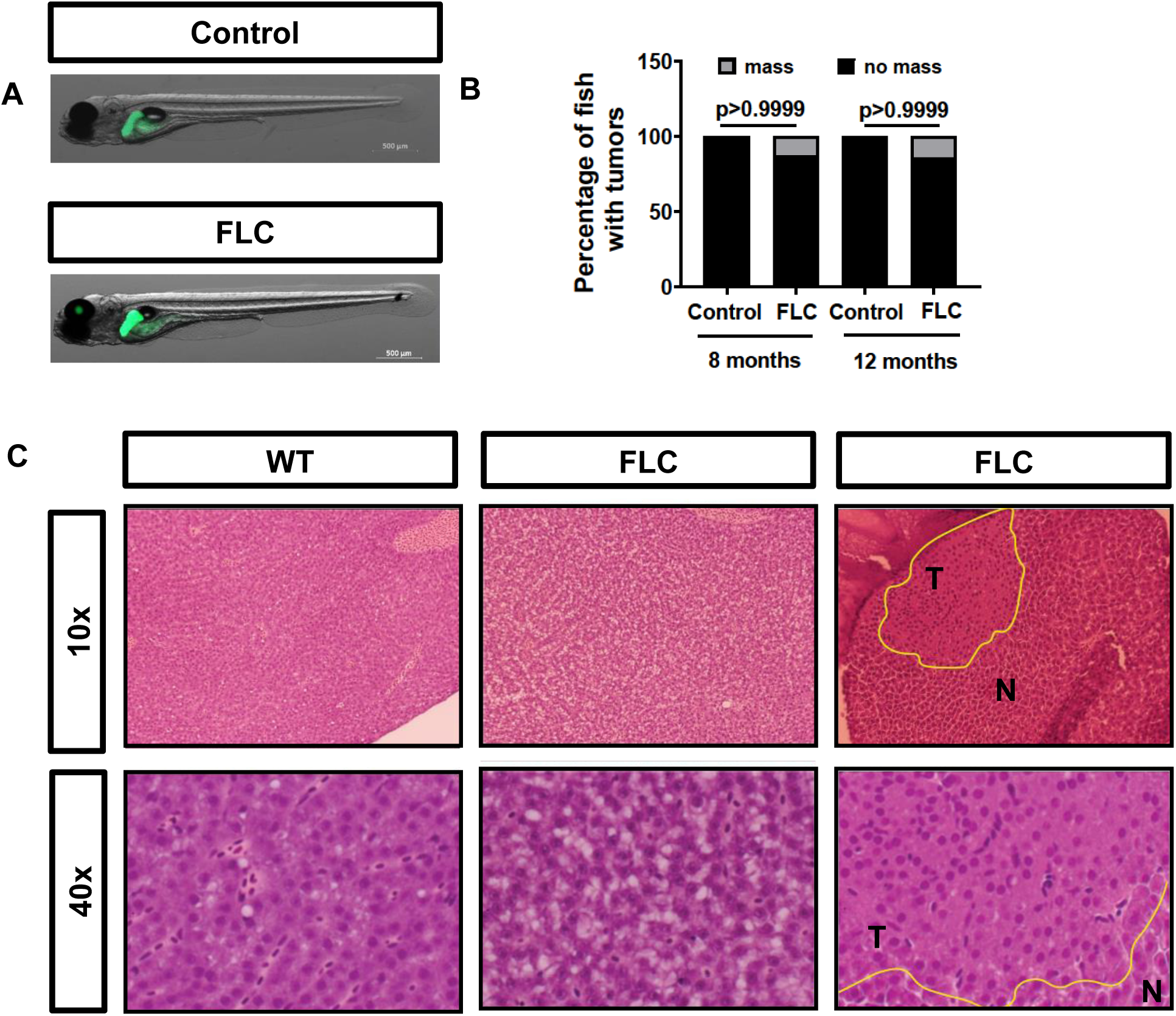
Overexpression of Dnajb1a-Prkacaa can induce liver masses at adult stages. **(A)** Representative maximum intensity projections of 7day post-fertilization (dpf) FLC larvae (*Tg(fabp10a:dnajb1a-prkacaa)/Tg(fabp10a:egfp-L10a)*) and Control siblings (*Tg(fabp10a:egfp-L10a)*); **(B)** Chi-square graphs showing percentage of larvae with masses; **(C)** Representative images of H&E staining of the livers of 12 month old fish at 10X and 40X magnification. Yellow outlines delineate masses.

**Table 1:** Zebrafish lines used in this study.

| Fish lines | ZFIN reference | Target | Genetic Background | Reference |
| --- | --- | --- | --- | --- |
| Casper | roy<sup>a9</sup>,<br>mitfa<sup>w2</sup> | - | Casper | White, RM<br>*et al.*,<br>2008 |
| *Tg(fabp10a: dnajb1a-prkacaa_cryaa:Cerulean)* | - | Hepatocytes<br>(screened for<br>eye marker) | Casper | This paper |
| *Tg(fabp10a: egfp-L10a)* | - | Hepatocyte<br>Ribosomes | Casper | This paper |
| *Tg(fabp10a: dnajb1a-prkacaa_cryaa:Cerulean/fabp10a:egfp-l10a)* | - | Hepatocytes | Casper | This paper |
| *Tg(fabp10a: mCherry-Kras)* | - | Hepatocytes<br>membrane | Casper | This paper |
| *Tg(mpeg1-mCherry-caax)* | - | Macrophages | Albinos | Bojarczuk,<br>A *et al.*;<br>2016 |
| *Tg(lyzc:bfp)* | - | Neutrophils | AB | De<br>Oliveira *et al.*, 2019 |
| *Tg(mpeg1-mCherry-caax/lyzc:bfp)* | - | Macrophages<br>and<br>Neutrophils | AB | This paper |
| *Tg(tnfa:gfp/mpeg1:mCherry-caax))* | - | Macrophages<br>and TNFa<br>expressing<br>cells/tissues | AB/Albinos | Miskolci, V<br>*et al.*,<br>2019 |

### FLC transgenic larvae display early malignancy features

Zebrafish larvae are a valuable model to study the early cellular and molecular events involved in liver cancer progression (de Oliveira et al., 2019; Huang et al., 2017; Nguyen et al., 2012; Yan et al., 2015; Yan et al., 2017; Zhao et al., 2016). Next, we wanted to determine if FLC transgenic larvae displayed early malignancy features. Liver size is a common measurement used to quantify liver disease progression (de Oliveira et al., 2019; Evason et al., 2015; Yan et al., 2015). Liver area and liver volume were increased in 7 days post-fertilization (dpf) FLC larvae compared to control siblings (Fig. 3A-C; Suppl. Movie S1 and S2). We next took advantage of the optical accessibility of zebrafish larvae to evaluate the size of hepatocytes *in vivo* by non-invasive live imaging. We outcrossed the FLC transgenic line, *Tg(fabp10a:dnajb1a-prkacaa_cryaa:Cerullean)*, with a line that expresses Kras in the hepatocyte membrane, *Tg(fabp10a:mCherry-Kras)* (Table 1). In FLC larvae, we observed an increase of hepatocytes area and diameter (Fig.3 D-F). Altogether, these data suggest that ectopic expression of *dnajb1a-prkacaa* fusion in hepatocytes induces hepatomegaly and that 7dpf FLC larvae can be used to study early FLC progression.

**Figure 3:**
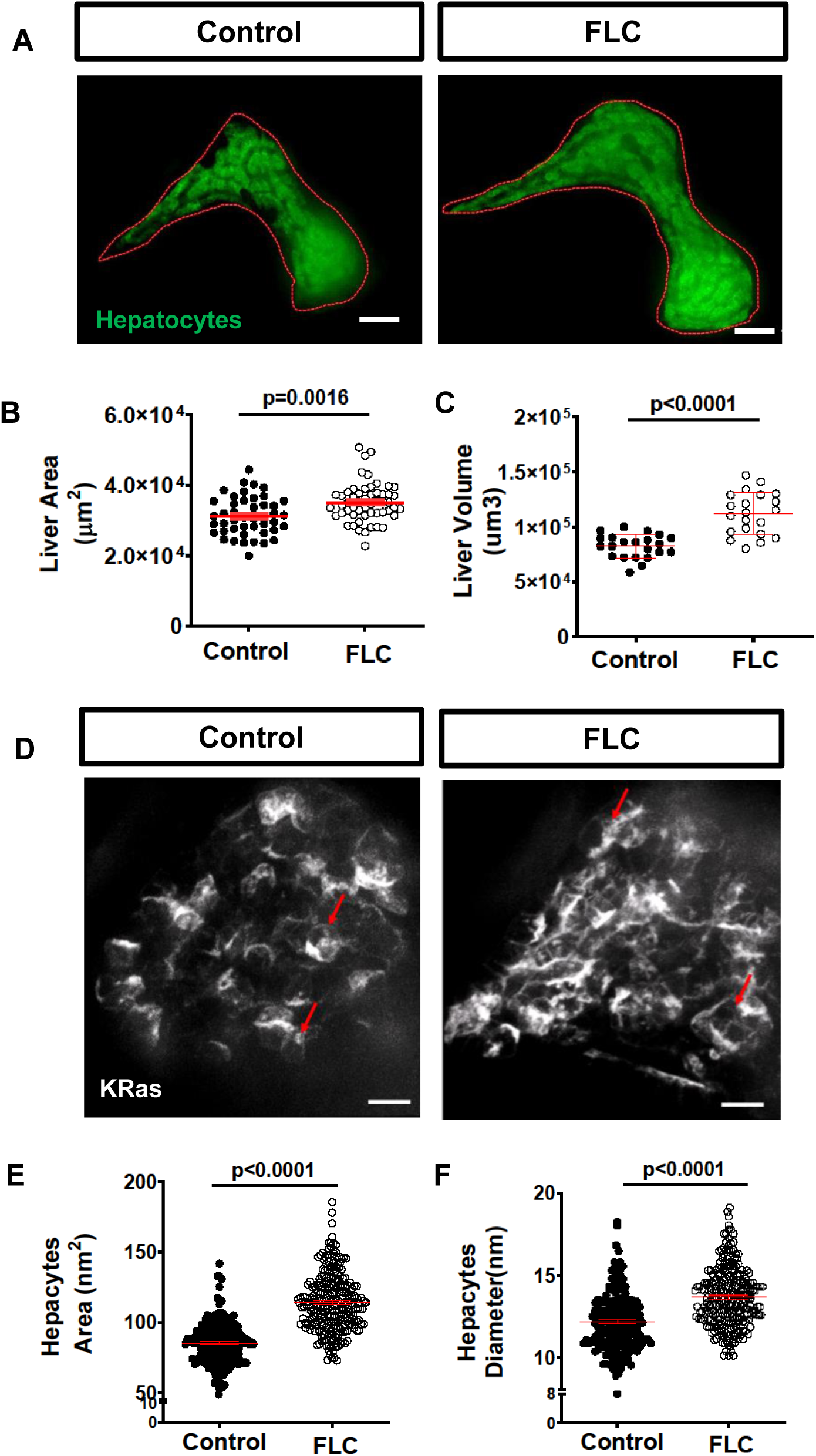
Overexpression of Dnajb1a-Prkacaa modulates liver morphology. **(A)** Representative maximum intensity projections of 7dpf FLC larvae (*Tg(fabp10a:dnajb1a-prkacaa)/Tg(fabp10a:egfp-L10a*)) and Control siblings (*Tg(fabp10a:egfp-L10a*)); **(B-C)** Graphs showing liver area (B) and liver volume (C). **(D)** Representative maximum intensity projections of 7dpf FLC larvae (*Tg(fabp10a:dnajb1a-prkacaa)/Tg(fabp10a:mCherryKras)*) and Control siblings (*Tg(fabp10a:mCherryKras)*).; **(E-F)** Graphs showing hepatocyte area (E) and diameter (F). Scale bar= 20μm. Data are from at least 3 independent experiments. Analysis performed in EEM in R. Dot-plots show mean ± SEM, p values are shown in each graph.

### Aberrant PKA induces innate immune cell infiltration in the liver area

Increased infiltration of neutrophils and macrophages has been associated with liver cancer progression (de Oliveira et al., 2019; Yan et al., 2015). It is still unclear how J-PKAca affects the immune cell composition in the liver microenvironment. We next outcrossed the FLC transgenic fish with labeled hepatocytes, *Tg(fabp10a:dnajb1a-prkacaa_cryaa:Cerullean)/(fabp10a:EGFP-l10),* with the double transgenic neutrophil and macrophage labeled line, *Tg(mpeg1-mCherry-caax/lyzc:bfp)* (Table 1) (Fig. 4A). We found that overexpression of the aberrant PKA increases both neutrophil and macrophage infiltration to the liver area in FLC transgenic larvae compared to control siblings (Fig. 4A-C). Time lapse-movies of the liver microenvironment area (Suppl. Movie S2) revealed robust recruitment of neutrophils to livers of FLC transgenic larvae compared to control siblings at this early phase. Livers of FLC larvae also exhibited an increased presence of macrophages in association with transformed hepatocytes that exhibited a round shape (Suppl. Movie S3 and S4). Overall, these data suggest that the presence of J-PKAca triggers an inflammatory response in the liver of FLC larvae.

**Figure 4:**
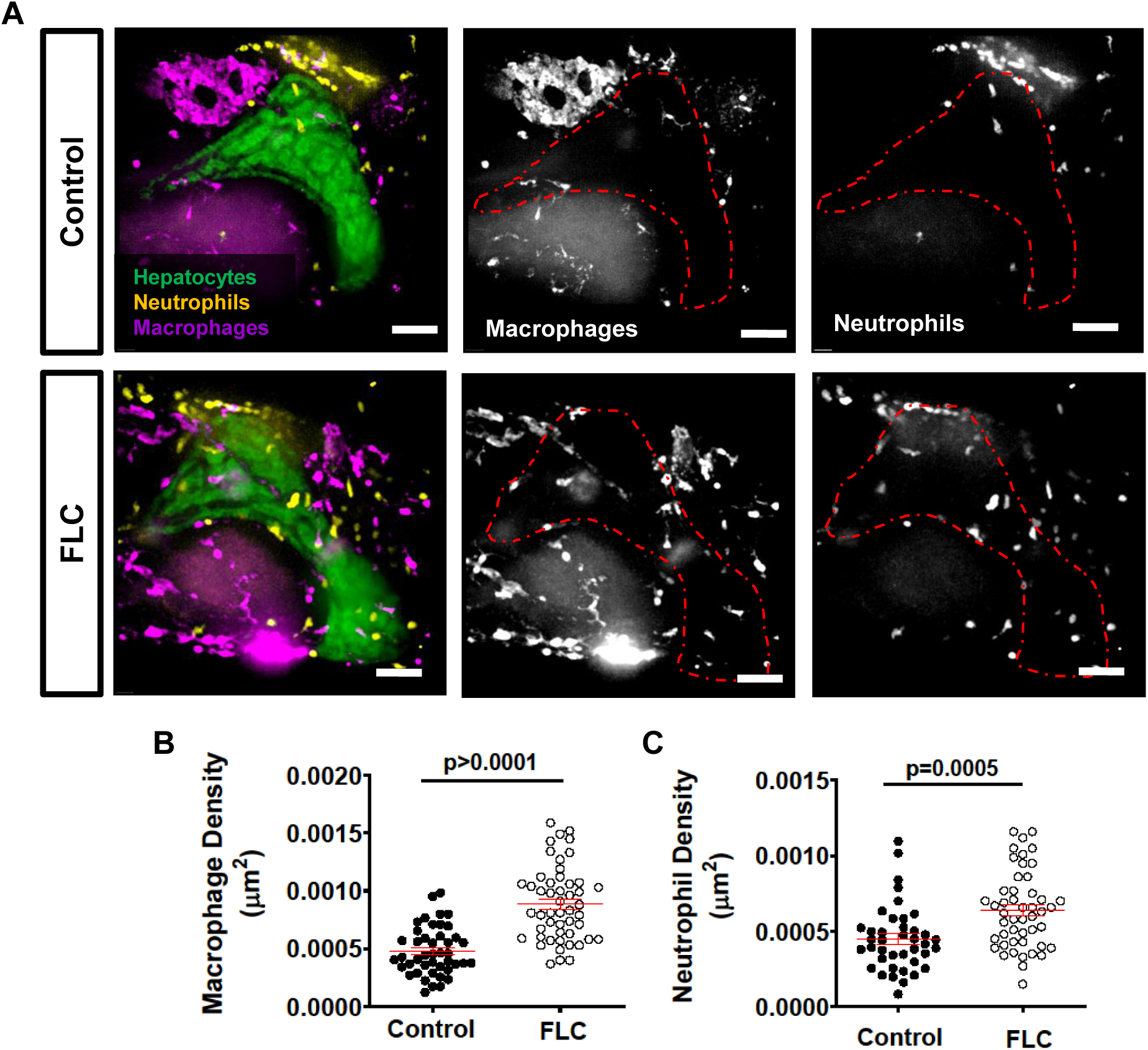
Overexpression of Dnajb1a-Prkacaa induces liver inflammation. **(A)** Representative maximum intensity projections of 7dpf FLC larvae (*Tg(fabp10a:dnajb1a-prkacaa)/Tg(fabp10a:egfp-L10a)/Tg(lyzC:bfp)/Tg(mpeg1:mCherry-caax)*) and Control siblings (*Tg(fabp10a:egfp-L10a)/Tg(lyzC:bfp)/Tg(mpeg1:mCherry-caax)*); **(B-C)** Graphs showing macrophage (B) and neutrophil (C) density at liver area. Scale bar= 40μm. Data are from at least from 3 independent experiments. Analysis performed in EEM in R. Dot-plots show mean ± SEM, p values are shown in each graph.

### FLC transgenic larvae have increased pro-inflammatory macrophages and Caspase-a activity in the liver

In our previous work, we found that the presence of pro-inflammatory macrophages in the liver microenvironment at early stages of progression in HCC larvae is associated with increased tumorigenesis in a model of non-alcoholic fatty liver disease-associated HCC (NAFLD-associated HCC) (de Oliveira et al., 2019). Nothing is known about the effect of J-PKAca on macrophage polarization *in vivo*. To identify pro-inflammatory macrophages, we outcrossed the FLC transgenic line, *Tg(fabp10a:dnajb1a-prkacaa_cryaa:Cerullean)*, with a reporter line of TNFα expression, *Tg(tnfa:egfp)* (Table 1). TNFα is a main molecular player in liver disease progression (Jang et al., 2014) and is mostly expressed by resident macrophages in the liver (Nakashima et al., 2013; Tosello-Trampont et al., 2012). We found that FLC transgenic larvae have increased numbers of TNFα-positive macrophages compared to control siblings (Fig. 4A-B). In a α recent study with a FLC murine model, a ssGSEA analysis for select functionally annotated gene sets showed an upregulation of genes associated with the inflammasome complex ((Kastenhuber et al., 2017). We therefore sought to determine if *dnajb1a-prkacaa* fusion transcript induces inflammasome activation via the activation of caspase-a, the zebrafish homologue for human caspase-1 (Angosto et al., 2012; Tyrkalska et al., 2016). Using FAM-FLICA assay, we found that caspase-a activity is significantly increased in the liver of FLC transgenic larvae compared to controls (Fig. 4C-D). Overall, our data suggest that J-PKAca promotes a pro-inflammatory liver microenvironment at early stages of FLC progression.

### Pharmacological inhibition of TNF-alpha secretion and caspase-a activity decreases inflammation and FLC progression

Zebrafish larvae are a powerful tool for small molecule screening in disease models(Wiley et al., 2017). Current therapeutic options for FLC patients are limited. Therefore, we decided to use a pharmacological approach to target inflammation and address its effect on FLC early progression. Our findings demonstrated an increase of pro-inflammatory macrophages (TNF-alpha positive) and caspase-a activity in the liver of FLC larvae (Fig. 4). Therefore, we next tested if inhibition of TNF-alpha secretion and caspase-a with pentoxifylline (PTX) and Ac-YVAD-CMK (C1INH), respectively, affected innate immune cell infiltration and liver size in FLC transgenic larvae. We observed that both treatments significantly decreased liver size as well as macrophage and neutrophil recruitment to the liver of FLC transgenic larvae (Fig. 5A-D). We also tested the effects of metformin on FLC. We and others have shown that metformin decreases inflammation and associated liver disease progression (de Oliveira et al., 2019; Li et al., 2019; Satapati et al., 2015). Surprisingly, we found that metformin treatment of FLC larvae did not affect liver size or macrophage infiltration (Fig. 5A-C). However, a small decrease in neutrophil infiltration was observed in FLC larvae treated with metformin (Fig. 5 A and D). Overall, our data suggest that TNF-alpha secretion and caspase-a activity mediate FLC-associated liver inflammation and may represent a new target to limit FLC progression.

**Figure 5:**
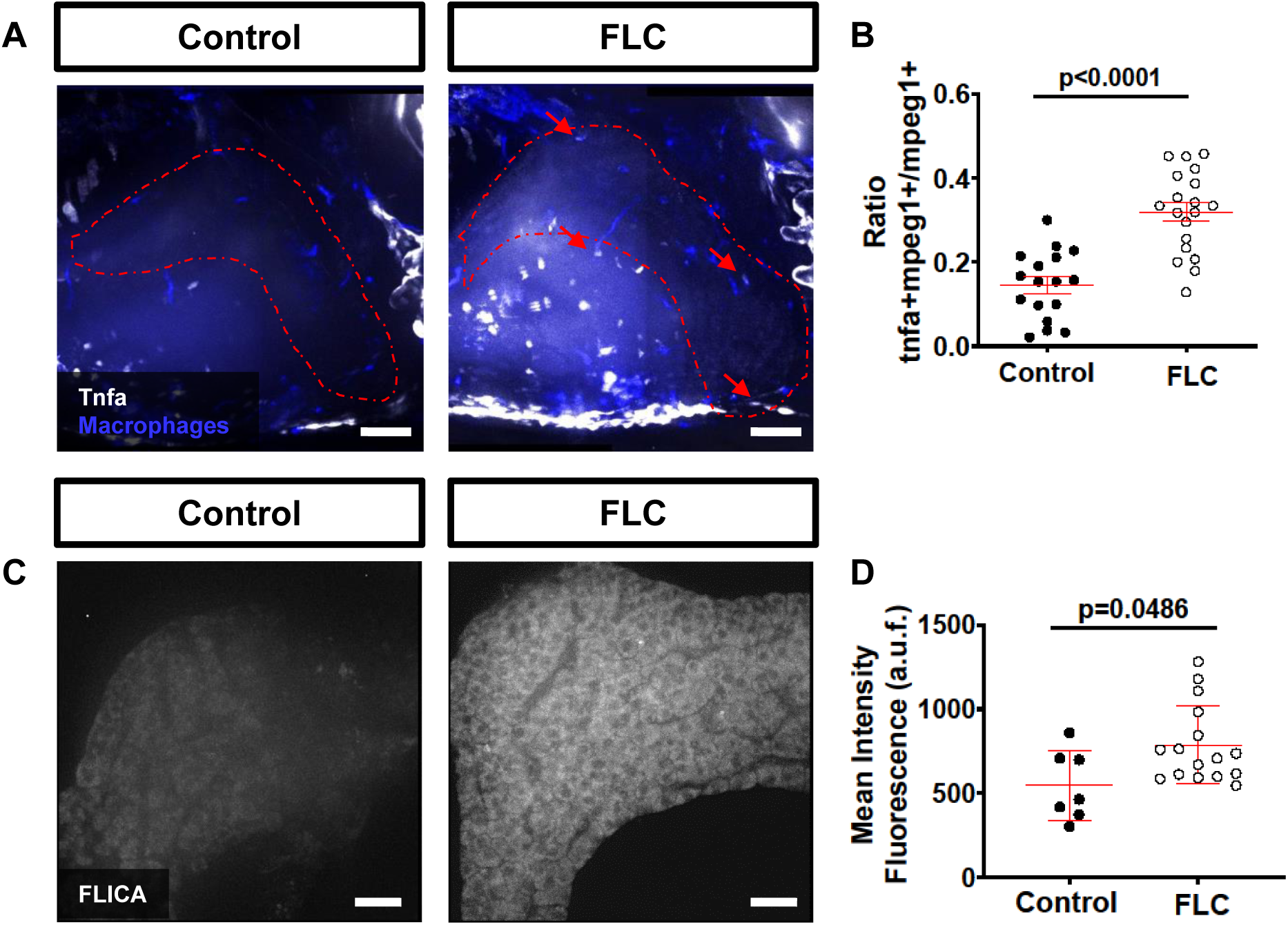
FLC larvae show increased pro-inflammatory macrophages and increased Caspase 1 activity at the liver area. **(A)** Representative maximum intensity projections of 7dpf FLC larvae (*Tg(fabp10a:dnajb1a-prkacaa_cryaa;Cerulean)/Tg(tnfa:gfp)/Tg(mpeg1:mCherry-caax*)) and Control siblings (*Tg(tnfa:gfp)/Tg(mpeg1:mCherry-caax*)); **(B)** Graph showing ratio of Tnfα positive macrophage over total macrophages at liver area **(C)** Representative maximum intensity projections of 7dpf FLC larvae (*Tg(fabp10a:dnajb1a-prkacaa_cryaa;Cerulean)* and Control wild type siblings; **(D)** Graph showing mean intensity fluorescent quantification at liver area; Scale bar=20μm. Data are from at least from 2 independent experiments. Analysis performed in EMM in R. Dot-plots show mean ± SEM, p values are shown in each graph. a.u.f= arbitrary units of fluorescence.

**Figure 6:**
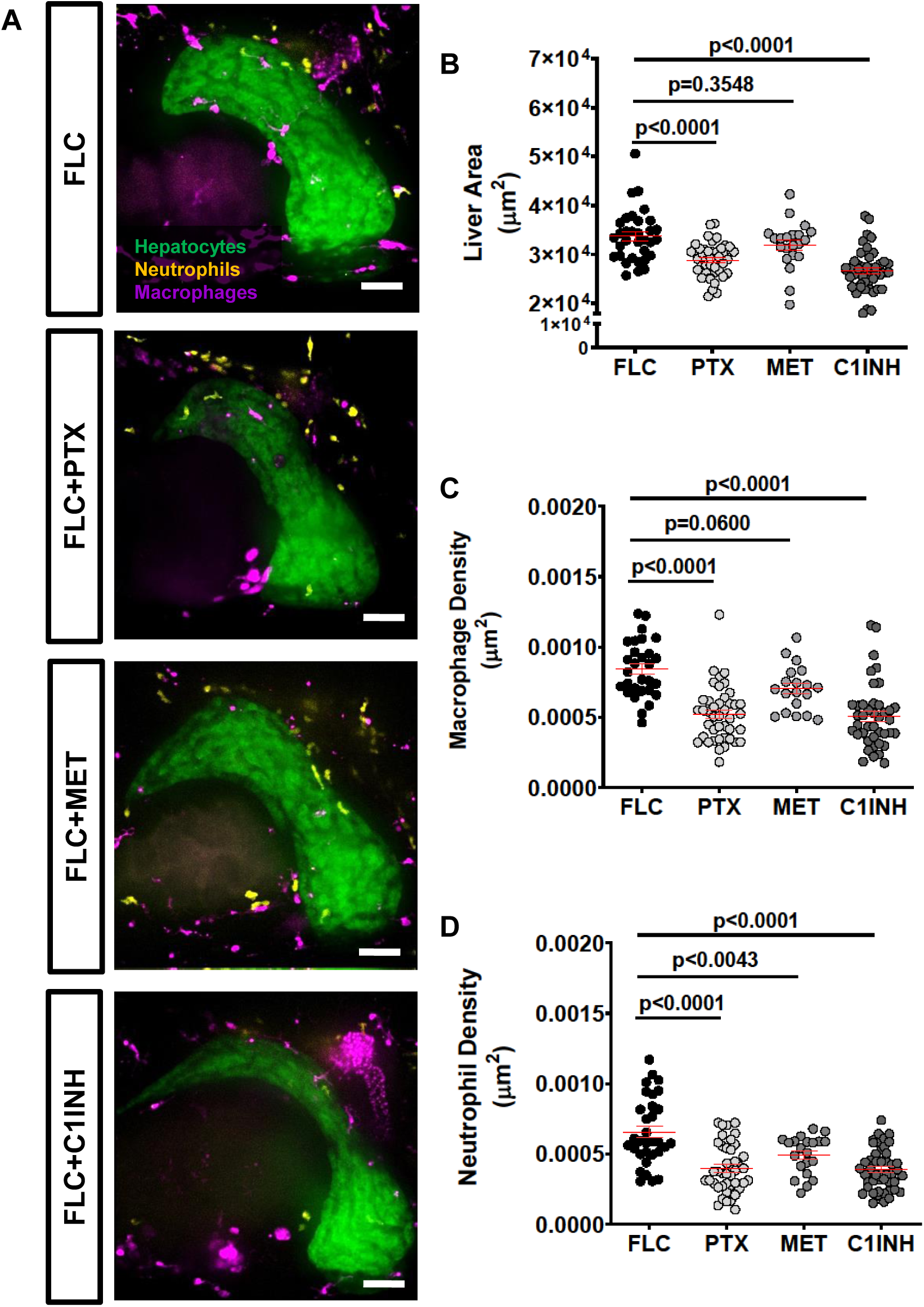
Pharmacological inhibition of TNF-alpha secretion and Caspase-a activity reduces inflammation and FLC progression. **(A)** Representative maximum intensity projections of 7dpf FLCa larvae (*Tg(fabp10a:dnajb1a-prkacaa)/Tg(fabp10a:egfp-L10a)/Tg(lyzC:bfp)/Tg(mpeg1:mCherry-caax*) and Control siblings (*Tg(fabp10a:egfp-L10a)/Tg(lyzC:bfp)/Tg(mpeg1:mCherry-caax)*) treated with 50μM Pentoxifylline (PTX), 50μM Metformin (MET) and 100 μM Ac-YVAD-CMK (C1INH). **(B-C)** Graphs showing liver area (B) macrophage and (C) neutrophil density. Scale bar= 40μm. Data are from at least 3 independent experiments. Analysis performed in EEM in R. Dot-plots show mean ± SEM, p values are shown in each graph.

## Discussion

Fibrolamellar Hepatocellular Carcinoma (FLC) is a rare pediatric liver cancer with few effective therapeutic options. The fusion chimera DNAJB1-PRKACA (J-PKAcα) has been identified as a unique driver of FLC (Graham et al., 2015; Honeyman et al., 2014; Simon et al., 2015). Here we report a zebrafish model for FLC generated by hepatocyte-specific ectopic expression of the zebrafish form of DNAJB1-PRKACA, *dnajab1a-prkacaa*. We find that the *dnajab1a-prkacaa* fusion transcript induces early hepatomegaly and inflammation in the liver area. One striking feature is that onset of inflammation is early and is characterized by the presence of TNFa positive macrophages, a feature that is not present in the standard catenin model of HCC (de Oliveira et al., 2019). In addition, this model provides a powerful tool to identify small molecules that alter inflammation and liver enlargement induced by *dnajab1a-prkacaa*.

Our findings show that the J-PKAcα fusion is sufficient to induce early stages of tumorigenesis, including liver enlargement and inflammation. However, there was a low incidence of mass formation in adult fish observed with *dnajab1a-prkacaa.* FLC transgenic fish develop masses, suggesting that, as in murine models, overexpression of *dnajab1a-prkacaa* is sufficient to drive tumorigenesis *in vivo*. However, FLC incidence was surprisingly low (1/8 at 8-months of age and 1/7 at 12-months). This may be due to low expression level or use of only one homologue of the DNAJB1 and PRKACA genes, the alpha form, *dnajab1a-prkacaa*, due to its higher identity and similarity with the human DNAJB1-PRKACA. However, the beta form *dnajb1b-prkacab* may have an important role in the liver acting in parallel with the alpha form and therefore affecting different pathways. In the future, it would be interesting to investigate whether the incidence of FLC tumorigenesis in zebrafish is increased by overexpression of *dnajb1b-prkacab* alone or combined with *dnajab1a-prkacaa*. In addition, as in the murine model, the masses found in FLC transgenic zebrafish lack markers of fibrosis seen in human FLC. This feature might be a crucial step for the progression of the disease and might be achieved with the use of fibrotic simulants (Kastenhuber et al., 2017). It is also possible that the activation of additional signaling pathways such as WNT/-β catenin (Kastenhuber et al., 2017) might be used to enhance *dnajb1a-prkacaa* tumorigenesis *in vivo* in zebrafish.

The function and activity of J-PKAcα has been another major focus of study in the field since drugs that target PKA activity could be a potential therapeutic option for FLC patients. Up to now, most data suggest that J-PKAcα fusion is enzymatically active and necessary for tumorigenesis *in vivo* (Kastenhuber et al., 2017). PKA activity mostly exerts an anti-inflammatory role (Campo et al., 2012) and is a major regulator of innate immune cells (Serezani et al., 2008). Several clinical drugs that target cAMP/PKA signaling pathway increase cAMP and are used to reduce inflammation and treat inflammatory disorders (Banner and Trevethick, 2004; Serezani et al., 2008). Importantly, immune cells are a main source of trophic support for transformed cancer cells and can play a role in the early progression of cancer (Feng et al., 2010; Giese et al., 2019; Powell and Huttenlocher, 2016) including liver cancer (Yan et al., 2015; Yan et al., 2017; Zhao et al., 2016). The role of J-PKAcα on the innate immune system is completely unclear. Using fluorescent labeled transgenic zebrafish larvae as a model, we and others are able to visualize the early interactions between immune cells and transformed hepatocytes and study the immune mechanisms involved in liver disease and cancer progression through non-invasive live imaging (de Oliveira et al., 2019; Yan et al., 2015; Yan et al., 2017). In theory, we would predict that the chimeric protein J-PKAcα would inhibit the innate immune response. Surprisingly, we observed the opposite-increased innate immune inflammation in our FLC transgenic model with increased neutrophil and macrophage infiltration into the liver, as well as.an increase of TNFα-positive macrophages. These data suggest that the J-PKAcα fusion protein promotes a pro-inflammatory liver microenvironment. The presence of uncontrolled fibrosis (Ward and Waxman, 2011), NF-κB activation (Li et al., 2010) and increased levels of CD68 (Ross et al., 2011), usually known as a cytoplasmic marker of macrophages and neutrophil granules (Ross et al., 2011), also suggests the presence of leukocytes in the liver microenvironment of FLC patients. Furthermore, dual oxidase 1 (DUOX1) is positively regulated by the cAMP/PKA cascade (Rigutto et al., 2009). DUOX1 is a main source of reactive oxygen species (ROS) production, such as hydrogen peroxide (H_2_O_2_), in epithelial tissues, including the liver. DUOX1 is overexpressed in liver tumors and further has been identified as a potential prognostic marker in HCC patients (Chen et al., 2016; Lu et al., 2011). H_2_O_2_ induces leukocyte recruitment and an inflammatory response after tissue damage (Candel et al., 2014; de Oliveira et al., 2015; de Oliveira et al., 2014; Niethammer et al., 2009; Razzell et al., 2013; Yoo et al., 2011) but also at early stages of cancer progression (Feng et al., 2010). Interestingly, murine and human FLC tumors show up-regulation of genes associated to ROS pathways, including enzymes involved in detoxifying ROS (Kastenhuber et al., 2017). It would be interesting to investigate if DUOX1/H_2_O_2_ signaling pathways are involved in the observed pro-inflammatory effect of J-PKAcα in our FLC zebrafish model. Moreover, we found increased caspase-a activity in the liver of FLC larvae, indicating increased activation of the inflammasome, in agreement with previous findings in human and murine FLC tumors (Kastenhuber et al., 2017).

Recurrence after complete surgical resection is common in FLC patients (Kassahun, 2016). The therapeutic strategies available for these patients with FLC relapse are limited and often not effective. Therefore, the discovery of new and improved therapeutic targets is a primary goal in this field. FLC zebrafish larvae models could be a powerful tool to aid in the identification of new drug targets using small molecule screening. Importantly, zebrafish larvae models display similar features of liver cancer progression (Goessling and Sadler, 2015; Huo et al., 2019; Lam et al., 2006; Li et al., 2014; Li et al., 2013; Wrighton et al., 2019; Zheng et al., 2013). Here we used a pharmacological approach to inhibit TNFα secretion and caspase-a activity in FLC larvae, since both TNFα and caspase activity were increased. We found that inhibition of both TNFα secretion and caspase-a activity both reduced innate immune cell infiltration in the liver as well as liver size, an established marker for liver disease progression (de Oliveira et al., 2019; Evason et al., 2015; Lam et al., 2006; Nguyen et al., 2012; Zheng et al., 2014). These findings support the potential of these approaches to reduce inflammation and decrease FLC progression. Further studies, with other FLC in vitro and *in vivo* models are needed to confirm the potential of these drugs to reduce FLC progression or recurrence.

Here we report a new FLC zebrafish model with unmatched non-invasive live imaging capabilities and scalability amenable to high-throughput drug screening. Overall, our findings support the idea that non-resolving inflammation might be fueling the liver microenvironment and contributing to FLC pathology. In addition, we found that pharmacological inhibition of TNFα secretion and caspase-a activity might be targets to treat inflammation and progression in FLC. In the future, it will be interesting to address how the FLC pro-inflammatory liver environment modulates the adaptive immune system; zebrafish models might be key tools in unraveling such mechanisms and finding new and improved therapeutic targets for FLC.

## Materials and Methods

### Zebrafish husbandry and maintenance

All protocols using zebrafish in this study were approved by the University of Wisconsin-Madison Institutional Animal Care and Use Committee. Adult zebrafish and embryos up to 5 days post-fertilization (dpf) were maintained as described previously (de Oliveira et al., 2019). At 5 dpf, larvae were transferred to 15cm petri dishes and kept in E3 media without methylene blue until 7dpf. For all experiments, larvae were anesthetized in E3 media without methylene blue containing 0.16 mg/ml Tricaine (MS222/ethyl 3-aminobenzoate; Sigma-Aldrich). Zebrafish lines used are summarized in Table 1.

#### Generation of Tg(fabp10a:dnajb1a-prkacaa_cryaa:Cerulean), Tg(fabp10a:egfp-l10a) and Tg(fabp10a:mcherry-Kras) lines

For *Tg(fabp10a:dnajb1a-prkacaa_cryaa:Cerulean),* DNA coding sequence for *dnajb1a-prkacaa* fusion gene was PCR amplified from a plasmid synthetized by IDT using following primers: Fw: 5’- CTTTGTGTTGATCGGGTACCGCCACCATGGGAAAAGATT -3’ Rv: 5’- CTGATTATGATCTAGACTAGAATTCAGCAAACTCCT -3’ Resulting PCR products were gel purified, and cloned using InFusion kit (Clontech) into an expression vector containing fabp10a promoter, minimal Tol2 elements for efficient integration and an SV40 polyadenylation sequence (Yoo et al., 2012), previously digested with KpnI and XbaI and gel purified. F0 Casper larvae were obtained by injecting 3 nL of 12.5 ng/mL DNA plasmid and 17.5 ng/mL in vitro transcribed (Ambion) transposase mRNA into the cell of one-cell stage embryo.

F0 larvae were raised to breeding age and crossed to adult Casper zebrafish. Founders were screened for Cerullean positive eye using a Zeiss Axio Zoom stereomicroscope (EMS3/SyCoP3; Zeiss; PlanNeoFluar Z 1X:0.25 FWD 56mm lens).

For *Tg(fabp10a:egfp-l10a)* DNA coding egfp-l10a was PCR amplified from a plasmid (Davenport et al., 2016), using following primers:

Fw: 5’- CTTTGTGTTGATCGggtaccGCCACCATGGTGAGCAAGGGCGAGGA -3’

Rv: 5’- CTGATTATGATCTAGACTAATACAGACGCTGGGGCTTGC -3’

Resulting PCR products were gel purified and cloned using InFusion kit (Clontech) into an expression vector containing fabp10a promoter sequence, minimal Tol2 elements for efficient integration and an SV40 polyadenylation sequence, previously digested with KpnI and XbaI and gel purified. Casper fish were injected and screened for EGFP expression in the liver as described above.

For *Tg(fabp10a:mCherry-Kras)* DNA coding Kras was PCR amplified from a plasmid (Freisinger and Huttenlocher, 2014), using following primers:

Fw: 5’- ACGAGCTGTACAAGTCCGGAATGACTGAATATAAACTTGTGGTGGTG -3’

Rv: 5’- CTGATTATGATCTAGATTACATAATTACACACTTTGTCTTTGACTTCT -3’

Resulting PCR products were gel purified and cloned using InFusion kit (Clontech) into an expression vector containing fabp10a promoter sequence, mCherry sequence, minimal Tol2 elements for efficient integration and an SV40 polyadenylation sequence, previously digested with BspEI and XbaI and gel purified. Casper fish were injected and screened for mCherry expression in the liver as described above.

### Liver dissection and Histology

Adult zebrafish (8 and 12 months of age) were euthanized by tricaine overdose and the livers removed by dissection. Livers were fixed in 10% formalin overnight. Samples were coded to facilitate blinded histopathologic evaluation. The livers were paraffin-embedded and 4um sections were prepared and stained with hematoxylin and eosin. Slides were evaluated by a board-certified veterinary pathologist (RH).

### Live imaging

All live imaging was performed using a zWEDGI device as previously described (Huemer et al., 2017). For time-lapse imaging, the loading chamber was filled with 1% low melting point agarose (Sigma) in E3 to retain the larvae in the proper position. Additional Tricaine/E3 was added as needed. All images were acquired with live larvae with the exception of FLICA staining. Images were acquired on a spinning disk confocal microscope (CSU-X; Yokogawa) with a confocal scanhead on a Zeiss Observer Z.1 inverted microscope equipped with a Photometrics Evolve EMCCD camera using a Plan-Apochromat 20x/0.8 M27 air objective with a 5 μm interval. For larvae with large livers, 2x2 tile images were taken.

### Liver and hepatocytes size measurements

For liver size measurements, 7dpf *Tg(fabp10a:dnajb1a-prkacaa_cryaa:Cerulean)* were outcrossed with *Tg*(*fabp10a:egfp-l10a*). For hepatocyte measurements, *Tg(fabp10a:dnajb1a-prkacaa_cryaa:Cerulean)* were outcrossed with *Tg(fabp10a:mCherry-Kras).* Liver area, liver volume, hepatocyte area and hepatocyte diameter were measured as previously described (de Oliveira et al., 2019).

### Quantification of neutrophil and macrophage recruitment

To quantify leukocyte recruitment we outcrossed double transgenic FLC line carrying the EGFP-L10a as a liver marker, *Tg(fabp10a:dnajb1a-prkacaa_cryaa:Cerulean)*/(*fabp10a:egfp-l10a*), with a double-transgenic line with labelled macrophages and neutrophils, *Tg(mpeg1-mCherry-caax/lyzc:bfp)*. After live imaging, Z series images were reconstructed in 2D maximum intensity projections (MIP) on ZEN pro 2012 software (Zeiss). Neutrophils, macrophages and counted within 50 μm from liver. Area of the liver was measured in each larva and used to normalize number of innate immune cells per liver area.

### TNFα-positive macrophages imaging and quantification

To assess TNFα-positive macrophages in the liver area we outcrossed the FLC line, *Tg(fabp10a:dnajb1a-prkacaa_cryaa:Cerulean),* with a double transgenic line labelled for macrophages and expressing EGFP under the TNFα promoter, (*Tg(mpeg1:mCherry-caax/TNFα:egfp))(Miskolci et al., 2019)*. After live imaging, Z series images were 3D reconstructed on Imaris software. Using the Imaris spots tool, total macrophages (mpeg1:mCherry-caax positive cells) were counted within 50μm from liver. TNFα positive macrophages (double positive mpeg1:mcherry/Tnfα:egfp cells) were quantified similarly. Z series images were reconstructed on Zen software to create maximum intensity projections.

### Drug treatment

Larvae were treated with metformin (MET), pentoxifylline (PTX) and Ac-YVAD-CMK (C1INH) as described previously (de Oliveira et al., 2019) (REF Sylwia paper). Briefly, we dissolved metformin (Enzo Life Sciences) in E3 without methylene blue at a final concentration of 50 µM. Pentoxifyline and AC-YVAD-CMK were first reconstituted in DMSO and later diluted 1000x in E3 without methylene blue at final concentration of 50 µM and 100 µM, respectively. Larvae were treated with these drugs from 3 to 7dpf. Drugs were freshly prepared and replaced daily.

### Caspase-1 activity assay

Larvae with 6dpf were incubated with FAM-FLICA (Immunochemistry Technologies) at 1:300 for 12h. Next day, larvae were washed with E3 and fixed in PIPES/1.5%Formaldehyde buffer at 4C. overnight. Larvae were washed 3 times with PBS 1x and livers were carefully dissected with the use of forceps on a stereomicroscope (Leica MZ 9.5). After dissection, livers were immersed in PBS and images were acquired on a spinning disk confocal microscope (CSU-X; Yokogawa) with a confocal scanhead on a Zeiss Observer Z.1 inverted microscope equipped with a Photometrics Evolve EMCCD camera using a EC Plan-Neofluor 40x/0.75 M27 air objective with a 1μm interval.

Mean Intensity fluorescence was measured on maximum intensity projections using Image J.

### Statistical analysis

All data plotted comprise at least three independent experimental replicates. Estimated Marginal Means (EMMs) analysis in R (www.r-project.org) (Vincent et al., 2016) was performed on pooled replicate experiments, using Tukey method when comparing more than two treatments. Graphical representations were done in GraphPad Prism version 6.

## Supporting information

Supplemental Movie S1 Control

Supplemental Movie S2 FLC

Supplemental Movie S3 Control

Supplemental Movie S4 FLC

## Acknowledgments

The authors would like to acknowledge to Dr. Leonard Zon, and Stephen Renshaw for *Tg(mpeg:mcherry-caax* line, and to Donghwan Jeon, Alyssa L. Graves and Leah Deshler for their technical support. We would like to acknowledge NIH/NCI R01 CA085862 to AH and funding from the Cancer Research Institute and Fibrolamellar Cancer Foundation to SdO.

## Notes

**Grant Support:** AH was funded by NCI CA085862; SdO is supported by Cancer Research Institute/Fibrolamellar Cancer Foundation

**Disclosures:** The authors disclose no conflicts.

## References

Angosto, D., Lopez-Castejon, G., Lopez-Munoz, A., Sepulcre, M. P., Arizcun, M., Meseguer, J. and Mulero, V. (2012). Evolution of inflammasome functions in vertebrates: Inflammasome and caspase-1 trigger fish macrophage cell death but are dispensable for the processing of IL-1beta. Innate Immun 18, 815–24.

Banner, K. H. and Trevethick, M. A. (2004). PDE4 inhibition: a novel approach for the treatment of inflammatory bowel disease. Trends in pharmacological sciences 25, 430–6.

Campo, G. M., Avenoso, A., D’Ascola, A., Prestipino, V., Scuruchi, M., Nastasi, G., Calatroni, A. and Campo, S. (2012). Protein kinase a mediated anti-inflammatory effects exerted by adenosine treatment in mouse chondrocytes stimulated with IL-1beta. Biofactors 38, 429–39.

Candel, S., de Oliveira, S., Lopez-Munoz, A., Garcia-Moreno, D., Espin-Palazon, R., Tyrkalska, S. D., Cayuela, M. L., Renshaw, S. A., Corbalan-Velez, R., Vidal-Abarca, I. et al. (2014). Tnfa signaling through tnfr2 protects skin against oxidative stress-induced inflammation. PLoS Biol 12, e1001855.

Chen, S., Ling, Q., Yu, K., Huang, C., Li, N., Zheng, J., Bao, S., Cheng, Q., Zhu, M. and Chen, M. (2016). Dual oxidase 1: A predictive tool for the prognosis of hepatocellular carcinoma patients. Oncol Rep 35, 3198–208.

Davenport, N. R., Sonnemann, K. J., Eliceiri, K. W. and Bement, W. M. (2016). Membrane dynamics during cellular wound repair. Mol Biol Cell 27, 2272–85.

de Oliveira, S., Boudinot, P., Calado, A. and Mulero, V. (2015). Duox1-derived H2O2 modulates Cxcl8 expression and neutrophil recruitment via JNK/c-JUN/AP-1 signaling and chromatin modifications. J Immunol 194, 1523–33.

de Oliveira, S., Houseright, R. A., Graves, A. L., Golenberg, N., Korte, B. G., Miskolci, V. and Huttenlocher, A. (2019). Metformin modulates innate immune-mediated inflammation and early progression of NAFLD-associated hepatocellular carcinoma in zebrafish. J Hepatol 70, 710–721.

de Oliveira, S., Lopez-Munoz, A., Candel, S., Pelegrin, P., Calado, A. and Mulero, V. (2014). ATP modulates acute inflammation in vivo through dual oxidase 1-derived H2O2 production and NF-kappaB activation. J Immunol 192, 5710–9.

Dhingra, S., Li, W., Tan, D., Zenali, M., Zhang, H. and Brown, R. E. (2010). Cell cycle biology of fibrolamellar hepatocellular carcinoma. International journal of clinical and experimental pathology 3, 792–7.

Engelholm, L. H., Riaz, A., Serra, D., Dagnaes-Hansen, F., Johansen, J. V., Santoni-Rugiu, E., Hansen, S. H., Niola, F. and Frodin, M. (2017). CRISPR/Cas9 Engineering of Adult Mouse Liver Demonstrates That the Dnajb1-Prkaca Gene Fusion Is Sufficient to Induce Tumors Resembling Fibrolamellar Hepatocellular Carcinoma. Gastroenterology 153, 1662–1673 e10.

Evason, K. J., Francisco, M. T., Juric, V., Balakrishnan, S., Lopez Pazmino Mdel, P., Gordan, J. D., Kakar, S., Spitsbergen, J., Goga, A. and Stainier, D. Y. (2015). Identification of Chemical Inhibitors of beta-Catenin-Driven Liver Tumorigenesis in Zebrafish. PLoS Genet 11, e1005305.

Feng, Y., Santoriello, C., Mione, M., Hurlstone, A. and Martin, P. (2010). Live imaging of innate immune cell sensing of transformed cells in zebrafish larvae: parallels between tumor initiation and wound inflammation. PLoS Biol 8, e1000562.

Freisinger, C. M. and Huttenlocher, A. (2014). Live imaging and gene expression analysis in zebrafish identifies a link between neutrophils and epithelial to mesenchymal transition. PLoS One 9, e112183.

Giese, M. A., Hind, L. E. and Huttenlocher, A. (2019). Neutrophil plasticity in the tumor microenvironment. Blood 133, 2159–2167.

Goessling, W. and Sadler, K. C. (2015). Zebrafish: an important tool for liver disease research. Gastroenterology 149, 1361–77.

Graham, R. P., Jin, L., Knutson, D. L., Kloft-Nelson, S. M., Greipp, P. T., Waldburger, N., Roessler, S., Longerich, T., Roberts, L. R., Oliveira, A. M. et al. (2015). DNAJB1-PRKACA is specific for fibrolamellar carcinoma. Mod Pathol 28, 822–9.

Graham, R. P., Lackner, C., Terracciano, L., Gonzalez-Cantu, Y., Maleszewski, J. J., Greipp, P. T., Simon, S. M. and Torbenson, M. S. (2018). Fibrolamellar carcinoma in the Carney complex: PRKAR1A loss instead of the classic DNAJB1-PRKACA fusion. Hepatology 68, 1441–1447.

Honeyman, J. N., Simon, E. P., Robine, N., Chiaroni-Clarke, R., Darcy, D. G., Lim, II, Gleason, C. E., Murphy, J. M., Rosenberg, B. R., Teegan, L. et al. (2014). Detection of a recurrent DNAJB1-PRKACA chimeric transcript in fibrolamellar hepatocellular carcinoma. Science 343, 1010–4.

Huang, S. J., Cheng, C. L., Chen, J. R., Gong, H. Y., Liu, W. and Wu, J. L. (2017). Inducible liver-specific overexpression of gankyrin in zebrafish results in spontaneous intrahepatic cholangiocarcinoma and hepatocellular carcinoma formation. Biochem Biophys Res Commun 490, 1052–1058.

Huemer, K., Squirrell, J. M., Swader, R., LeBert, D. C., Huttenlocher, A. and Eliceiri, K. W. (2017). zWEDGI: Wounding and Entrapment Device for Imaging Live Zebrafish Larvae. Zebrafish 14, 42–50.

Huo, X., Li, H., Li, Z., Yan, C., Mathavan, S., Liu, J. and Gong, Z. (2019). Transcriptomic analyses of oncogenic hepatocytes reveal common and different molecular pathways of hepatocarcinogenesis in different developmental stages and genders in kras(G12V) transgenic zebrafish. Biochem Biophys Res Commun 510, 558–564.

Jang, M. K., Kim, H. S. and Chung, Y. H. (2014). Clinical aspects of tumor necrosis factor-alpha signaling in hepatocellular carcinoma. Curr Pharm Des 20, 2799–808.

Kassahun, W. T. (2016). Contemporary management of fibrolamellar hepatocellular carcinoma: diagnosis, treatment, outcome, prognostic factors, and recent developments. World J Surg Oncol 14, 151.

Kastenhuber, E. R., Lalazar, G., Houlihan, S. L., Tschaharganeh, D. F., Baslan, T., Chen, C. C., Requena, D., Tian, S., Bosbach, B., Wilkinson, J. E. et al. (2017). DNAJB1-PRKACA fusion kinase interacts with beta-catenin and the liver regenerative response to drive fibrolamellar hepatocellular carcinoma. Proc Natl Acad Sci U S A 114, 13076–13084.

Kuang, D. M., Zhao, Q., Peng, C., Xu, J., Zhang, J. P., Wu, C. and Zheng, L. (2009). Activated monocytes in peritumoral stroma of hepatocellular carcinoma foster immune privilege and disease progression through PD-L1. J Exp Med 206, 1327–37.

Kuang, D. M., Zhao, Q., Wu, Y., Peng, C., Wang, J., Xu, Z., Yin, X. Y. and Zheng, L. (2011). Peritumoral neutrophils link inflammatory response to disease progression by fostering angiogenesis in hepatocellular carcinoma. J Hepatol 54, 948–55.

Lam, S. H., Wu, Y. L., Vega, V. B., Miller, L. D., Spitsbergen, J., Tong, Y., Zhan, H., Govindarajan, K. R., Lee, S., Mathavan, S. et al. (2006). Conservation of gene expression signatures between zebrafish and human liver tumors and tumor progression. Nat Biotechnol 24, 73–5.

Li, W., Tan, D., Zenali, M. J. and Brown, R. E. (2010). Constitutive activation of nuclear factor-kappa B (NF-kB) signaling pathway in fibrolamellar hepatocellular carcinoma. International journal of clinical and experimental pathology 3, 238–43.

Li, X. F., Chen, D. P., Ouyang, F. Z., Chen, M. M., Wu, Y., Kuang, D. M. and Zheng, L. (2015). Increased autophagy sustains the survival and pro-tumourigenic effects of neutrophils in human hepatocellular carcinoma. Journal of hepatology 62, 131–9.

Li, Y. L., Li, X. Q., Wang, Y. D., Shen, C. and Zhao, C. Y. (2019). Metformin alleviates inflammatory response in non-alcoholic steatohepatitis by restraining signal transducer and activator of transcription 3-mediated autophagy inhibition in vitro and in vivo. Biochem Biophys Res Commun 513, 64–72.

Li, Z., Luo, H., Li, C., Huo, X., Yan, C., Huang, X., Al-Haddawi, M., Mathavan, S. and Gong, Z. (2014). Transcriptomic analysis of a transgenic zebrafish hepatocellular carcinoma model reveals a prominent role of immune responses in tumour progression and regression. International journal of cancer. Journal international du cancer 135, 1564–73.

Li, Z., Zheng, W., Wang, Z., Zeng, Z., Zhan, H., Li, C., Zhou, L., Yan, C., Spitsbergen, J. M. and Gong, Z. (2013). A transgenic zebrafish liver tumor model with inducible Myc expression reveals conserved Myc signatures with mammalian liver tumors. Dis Model Mech 6, 414–23.

Lu, C. L., Qiu, J. L., Huang, P. Z., Zou, R. H., Hong, J., Li, B. K., Chen, G. H. and Yuan, Y. F. (2011). NADPH oxidase DUOX1 and DUOX2 but not NOX4 are independent predictors in hepatocellular carcinoma after hepatectomy. Tumour Biol 32, 1173–82.

Miskolci, V., Squirrell, J., Rindy, J., Vincent, W., Sauer, J. D., Gibson, A., Eliceiri, K. W. and Huttenlocher, A. (2019). Distinct inflammatory and wound healing responses to complex caudal fin injuries of larval zebrafish. Elife 8.

Nakashima, H., Ogawa, Y., Shono, S., Kinoshita, M., Nakashima, M., Sato, A., Ikarashi, M. and Seki, S. (2013). Activation of CD11b+ Kupffer cells/macrophages as a common cause for exacerbation of TNF/Fas-ligand-dependent hepatitis in hypercholesterolemic mice. PLoS One 8, e49339.

Nguyen, A. T., Emelyanov, A., Koh, C. H., Spitsbergen, J. M., Parinov, S. and Gong, Z. (2012). An inducible kras(V12) transgenic zebrafish model for liver tumorigenesis and chemical drug screening. Dis Model Mech 5, 63–72.

Niethammer, P., Grabher, C., Look, A. T. and Mitchison, T. J. (2009). A tissue-scale gradient of hydrogen peroxide mediates rapid wound detection in zebrafish. Nature 459, 996–9.

Oikawa, T., Wauthier, E., Dinh, T. A., Selitsky, S. R., Reyna-Neyra, A., Carpino, G., Levine, R., Cardinale, V., Klimstra, D., Gaudio, E. et al. (2015). Model of fibrolamellar hepatocellular carcinomas reveals striking enrichment in cancer stem cells. Nat Commun 6, 8070.

Powell, D. R. and Huttenlocher, A. (2016). Neutrophils in the Tumor Microenvironment. Trends in immunology 37, 41–52.

Razzell, W., Evans, I. R., Martin, P. and Wood, W. (2013). Calcium flashes orchestrate the wound inflammatory response through DUOX activation and hydrogen peroxide release. Curr Biol 23, 424–9.

Riggle, K. M., Riehle, K. J., Kenerson, H. L., Turnham, R., Homma, M. K., Kazami, M., Samelson, B., Bauer, R., McKnight, G. S., Scott, J. D. et al. (2016). Enhanced cAMP-stimulated protein kinase A activity in human fibrolamellar hepatocellular carcinoma. Pediatric research 80, 110–8.

Rigutto, S., Hoste, C., Grasberger, H., Milenkovic, M., Communi, D., Dumont, J. E., Corvilain, B., Miot, F. and De Deken, X. (2009). Activation of dual oxidases Duox1 and Duox2: differential regulation mediated by camp-dependent protein kinase and protein kinase C-dependent phosphorylation. J Biol Chem 284, 6725–34.

Ross, H. M., Daniel, H. D., Vivekanandan, P., Kannangai, R., Yeh, M. M., Wu, T. T., Makhlouf, H. R. and Torbenson, M. (2011). Fibrolamellar carcinomas are positive for CD68. Modern pathology: an official journal of the United States and Canadian Academy of Pathology, Inc 24, 390–5.

Satapati, S., Kucejova, B., Duarte, J. A., Fletcher, J. A., Reynolds, L., Sunny, N. E., He, T., Nair, L. A., Livingston, K. A., Fu, X. et al. (2015). Mitochondrial metabolism mediates oxidative stress and inflammation in fatty liver. J Clin Invest 125, 4447–62.

Serezani, C. H., Ballinger, M. N., Aronoff, D. M. and Peters-Golden, M. (2008). Cyclic AMP: master regulator of innate immune cell function. American journal of respiratory cell and molecular biology 39, 127–32.

Simon, E. P., Freije, C. A., Farber, B. A., Lalazar, G., Darcy, D. G., Honeyman, J. N., Chiaroni-Clarke, R., Dill, B. D., Molina, H., Bhanot, U. K. et al. (2015). Transcriptomic characterization of fibrolamellar hepatocellular carcinoma. Proceedings of the National Academy of Sciences of the United States of America 112, E5916–25.

Skalhegg, B. S., Funderud, A., Henanger, H. H., Hafte, T. T., Larsen, A. C., Kvissel, A. K., Eikvar, S. and Orstavik, S. (2005). Protein kinase A (PKA)--a potential target for therapeutic intervention of dysfunctional immune cells. Current drug targets 6, 655–64.

Tosello-Trampont, A. C., Landes, S. G., Nguyen, V., Novobrantseva, T. I. and Hahn, Y. S. (2012). Kuppfer cells trigger nonalcoholic steatohepatitis development in diet-induced mouse model through tumor necrosis factor-alpha production. J Biol Chem 287, 40161–72.

Tyrkalska, S. D., Candel, S., Angosto, D., Gomez-Abellan, V., Martin-Sanchez, F., Garcia-Moreno, D., Zapata-Perez, R., Sanchez-Ferrer, A., Sepulcre, M. P., Pelegrin, P. et al. (2016). Neutrophils mediate Salmonella Typhimurium clearance through the GBP4 inflammasome-dependent production of prostaglandins. Nat Commun 7, 12077.

Vincent, W. J., Freisinger, C. M., Lam, P. Y., Huttenlocher, A. and Sauer, J. D. (2016). Macrophages mediate flagellin induced inflammasome activation and host defense in zebrafish. Cell Microbiol 18, 591–604.

Ward, S. C. and Waxman, S. (2011). Fibrolamellar carcinoma: a review with focus on genetics and comparison to other malignant primary liver tumors. Semin Liver Dis 31, 61–70.

Wiley, D. S., Redfield, S. E. and Zon, L. I. (2017). Chemical screening in zebrafish for novel biological and therapeutic discovery. Methods Cell Biol 138, 651–679.

Wrighton, P. J., Oderberg, I. M. and Goessling, W. (2019). There Is Something Fishy About Liver Cancer: Zebrafish Models of Hepatocellular Carcinoma. Cell Mol Gastroenterol Hepatol 8, 347–363.

Xu, L., Hazard, F. K., Zmoos, A. F., Jahchan, N., Chaib, H., Garfin, P. M., Rangaswami, A., Snyder, M. P. and Sage, J. (2015). Genomic analysis of fibrolamellar hepatocellular carcinoma. Hum Mol Genet 24, 50–63.

Yan, C., Huo, X., Wang, S., Feng, Y. and Gong, Z. (2015). Stimulation of hepatocarcinogenesis by neutrophils upon induction of oncogenic kras expression in transgenic zebrafish. J Hepatol 63, 420–8.

Yan, C., Yang, Q. and Gong, Z. (2017). Tumor-Associated Neutrophils and Macrophages Promote Gender Disparity in Hepatocellular Carcinoma in Zebrafish. Cancer Res 77, 1395–1407.

Yoo, S. K., Lam, P. Y., Eichelberg, M. R., Zasadil, L., Bement, W. M. and Huttenlocher, A. (2012). The role of microtubules in neutrophil polarity and migration in live zebrafish. J Cell Sci 125, 5702–10.

Yoo, S. K., Starnes, T. W., Deng, Q. and Huttenlocher, A. (2011). Lyn is a redox sensor that mediates leukocyte wound attraction in vivo. Nature 480, 109–12.

Zhao, Y., Huang, X., Ding, T. W. and Gong, Z. (2016). Enhanced angiogenesis, hypoxia and neutrophil recruitment during Myc-induced liver tumorigenesis in zebrafish. Scientific reports 6, 31952.

Zheng, W., Li, Z., Nguyen, A. T., Li, C., Emelyanov, A. and Gong, Z. (2014). Xmrk, kras and myc transgenic zebrafish liver cancer models share molecular signatures with subsets of human hepatocellular carcinoma. PLoS One 9, e91179.

Zheng, W., Xu, H., Lam, S. H., Luo, H., Karuturi, R. K. and Gong, Z. (2013). Transcriptomic analyses of sexual dimorphism of the zebrafish liver and the effect of sex hormones. PLoS One 8, e53562.

